# The interaction of orientation-specific surround suppression and visual-spatial attention

**DOI:** 10.1101/091553

**Authors:** Ariel Rokem, Ayelet Nina Landau

## Abstract

Orientation selective surround suppression (OSSS) is a reduction in the perceived contrast of a stimulus, which occurs when a collinear grating is placed adjacent to the stimulus. Attention affects performance on many visual tasks, and we asked whether the perceptual effects of OSSS are mitigated through the allocation of voluntary visual-spatial attention. Participants were tested in a contrast discrimination task: at the beginning of each trial, one location on the screen was cued and a subsequent contrast judgment was then more likely (70%) to be performed in that location. Replicating previous results, we found that the point of subjective equality (PSE) was elevated for a collinear, relative to an orthogonal, surround. While the PSE was similar for validly and invalidly cued trials, the just noticeable difference (JND) was larger for invalid cue trials, and for collinear, relative to orthogonal surround, suggesting that while OSSS affects both perceived contrast and sensitivity, voluntary attention affects only perceptual sensitivity. In another experiment no informative cue was provided, and attention was distributed over the entire display. In this case, JND and PSE were shifted depending on the contrast of the distractor, suggesting that OSSS is affected by the allocation of visual-spatial attention, but only under conditions of distributed attention.

## Introduction

Visual perception is subject to modulation by both bottom-up effects, such as the configural juxtaposition of different stimuli, and top-down effects such as attention. In the present study we investigated the interaction between a top-down effect, the allocation of voluntary visual spatial attention, and a bottom-up effect, the suppression of contrast perception by surrounding stimuli. Specifically, we asked how contrast perception is modulated by these two factors, and by their combination.

### Voluntary visual-spatial attention affects perception

Visual-spatial attention improves our ability to extract useful information from visual scenes, by selecting certain aspects of the scene and enhancing sensitivity to these aspects. Perception is therefore affected by expectations about currently relevant aspects of a visual scene. For example, information indicating that part of the visual field is relevant to our current goals may affect the manner in which visual information at that location is processed, relative to other irrelevant locations. Voluntary visual-spatial attention affects performance on many different perceptual tasks, including sensitivity (Bashinski & Bacharach, 1980) and response time (Posner, Snyder, & Davidson, 1980) to high-contrast stimuli, the segmentation of visual texture stimuli (Yeshurun, Montagna, & Carrasco, 2008), and the detection of low-contrast gratings embedded in noise (Dosher & Lu, 2000), or masked with a backward mask (Smith, 2000). However, whether attention affects the appearance of the discriminated stimuli is a matter of an ongoing debate (Abrams, Barbot, & Carrasco, 2010; Anton-erxleben, Abrams, & Carrasco, 2010; Carrasco, Ling, & Read, 2004a; Ling & Carrasco, 2007; Liu, Abrams, & Carrasco, 2009; Schneider, 2008, 2011). In the present study, we investigated whether information about the location of a task-relevant stimulus would affect sensitivity, appearance, or both.

No stimulus is an island. Natural stimuli often appear in the context of visual scenes that contain multiple other stimuli. The modulation of perception through spatial context appears in different guises (Albright & Stoner, 2002; Graham, 2011), and may underlie important perceptual functions, such as the recovery of depth information, scene segregation and the maintenance of perceptual constancies. Previous studies have examined the effects of allocation of attention on contextual effects, such as adaptation (Pestilli & Carrasco, 2007) and crowding (Yeshurun & Rashal, 2010). Meanwhile, neurophysiological studies have demonstrated that visual-spatial attention may also alleviate the effects of surround suppression (Sundberg, Mitchell, & Reynolds, 2009). To bridge between these neurophysiological findings and the previous behavioral findings, the present study examined the manner in which the allocation of voluntary attention affects the perception of surround-suppressed stimuli.

### Orientation selective surround suppression (OSSS)

One of the widely studied contextual effects on visual perception is the reduction in the perceived contrast of a stimulus, when it is adjacent to a stimulus of higher contrast. This phenomenon, known as surround suppression (because it has most often been studied using a high contrast stimulus that surrounds a low contrast stimulus, but see Petrov and McKee 2006) has been observed in different stimuli, including random textures (Chubb, Sperling, & Solomon, 1989; Dakin, Carlin, & Hemsley, 2005) plaids (Olzak & Laurinen, 1999), and oriented gratings (Ejima & Takahashi, 1985; Solomon, Sperling, & Chubb, 1993, Cannon & Fullenkamp, 1991; Xing & Heeger, 2000, 2001; Yu, Klein, & Levi, 2001). In studies using grating stimuli, surround suppression has been found to be more powerful when the surrounding and surrounded stimuli have collinear orientations, relative to when they have different orientations. Therefore, recent studies have often focused on the orientation-selective component of surround suppression (OSSS), by comparing suppression for parallel and orthogonal gratings on a within-subject basis (Kosovicheva, Sheremata, Rokem, Landau, & Silver, 2012; Yoon et al., 2009; Yoon et al., 2010). This form of suppression is also the focus of the present study.

To study the interaction of attention and OSSS, we combined a contrast-matching task (Yu et al., 2001; Yu & Levi, 2000) within the spatial cuing paradigm (Posner, 1980). In the contrast-matching task, participants were asked to choose which of two intervals contained a higher contrast stimulus. The first interval always contained two grating stimuli that were surrounded by grating annuli; the surround stimuli were either orthogonally or collinearly oriented with the center grating stimuli. The second interval contained a single grating presented in isolation at one of the previously presented gratings location (Figure 1). Participants were instructed to compare the contrast of the gratings in the two intervals that were co-localized on the screen. Additionally, trials were preceded by an informative cue that indicated the likely location of the grating to be discriminated.

We found that visual-spatial attention affected sensitivity (i.e. the ability to discriminate fine changes in contrast), but it did not affect the decrease in perceived contrast that resulted from OSSS. This suggests that the bias in the appearance of the stimulus, induced by OSSS, and sensitivity to contrast changes, modulated by attention, are governed by different mechanisms. As in the case of adaptation (Pestilli & Carrasco, 2007), voluntary visual-spatial attention seems to operate on the already suppressed stimulus, at a stage subsequent to the suppressive effect itself.

### Focused versus distributed attention and the effects of involuntary attention

The spatial distribution of visual-spatial attention in the visual field also affects the way in which attention and perception interact (Herrmann, Montaser-Kouhsari, Carrasco, & Heeger, 2010; Reynolds & Heeger, 2009; Schwartz & Coen-cagli, 2013). For example, thresholds in a contrast discrimination task were elevated in the presence of high contrast distractors (Pestilli, Carrasco, Heeger, & Gardner, 2011). This suggests that contrast discrimination is affected by elements of the display that are not directly relevant to current task performance. This is because in addition to the voluntary allocation of attention to a specific location, salient stimuli in the visual field (e.g. high-contrast distractor stimuli) can elicit involuntary attention (Prinzmetal & Landau, 2007). The perceptual and behavioral effects of involuntary attention are distinctly different from the effects of voluntary attention (Prinzmetal, Mccool, & Park, 2005; Prinzmetal, Zvinyatskovskiy, Gutierrez, & Dilem, 2008), as are the neural mechanisms underlying these two forms of attention (Esterman et al., 2008; Landau, Esterman, Robertson, Bentin, & Prinzmetal, 2007; Rokem, Landau, Garg, Prinzmetal, & Silver, 2010). Importantly, the degree to which involuntary attention affects performance, e.g. the effects of distractor stimuli, may depend on the demands of the task performed: for example, the elevation in contrast discrimination thresholds was larger when attention was distributed over the visual field, relative to when attention was selectively focused in one location (Pestilli et al., 2011). These differences in task effects in focused and distributed attention are evidence of the different fate of the distractors under different conditions: when attention is focused in a narrow part of the visual field, blocking of other stimuli is possible. On the other hand, when attention is distributed more widely, stimuli from other parts of the visual field are merely attenuated (Yigit-Elliott, Palmer, & Moore, 2011).

To test the effects of the spatial distribution of attention, we compared task performance in the focused attention (informative cues) condition to performance in another OSSS experiment, in which uninformative (neutral) cues were presented. The neutral cues contained the same timing information as the informative cues, but no spatial information about the location in which contrast discrimination would be performed (Figure 1). We found that when the cues contained no spatial information and attention was spatially distributed, the contrast of the other stimulus (the distractor) substantially modulated perception: bias due to OSSS was increased in the presence of a high contrast distractor and decreased in the presence of a low contrast distractor. In addition, sensitivity decreased due to involuntary allocation of attention to the distractor, in a manner similar to the effects of voluntary allocation of attention away from the target in focused attention conditions.

## Methods

### Subjects

16 healthy subjects participated in the experiment (Age: 27.5 +/−4; 11 female). All participants had previous experience of participation in psychophysical tasks. Participants provided informed consent and the study protocol was approved by the ethics committee of the Medical school at the Johann Wolfgang Goethe-University, Frankfurt.

### Stimuli and procedure

Stimuli were presented on a 22'' Samsung 2233RZ LCD display (Wang and Nikolic, 2011). Gray levels were measured with a photometer and corrected to be linear. Stimuli were presented using software written with Psychopy (Peirce, 2007). Stimuli were gray-scale vertical or horizontal sinusoidal grating patches presented centered at 6 degrees of visual angle eccentric to fixation, on either side of the fixation point (Figure 1). The spatial frequency of the gratings was 2 cycles/degree and their size was 3 degrees visual angle (dva) in diameter. The maximal luminance of the display was 338 cd/m^2^. In each trial, participants were presented with gratings on two intervals (Figure 1C and 1E). In the first interval two gratings were presented on either side of the fixation point (Figure 1C). These gratings could have one of the following contrast levels: 1%, 10%, 20%, 30%, 40%, 50%, 55%, 60%, 65%, 70%, or 90%. We refer to this contrast as the *comparison stimulus.* They were surrounded by other full contrast (100% contrast) grating annuli extending 8 dva in diameter. A thin (0.15 dva) black dividing line separated the central grating patches and surround annulus stimuli. Surround annulus stimuli were either oriented at the same orientation as the central grating patches, or oriented orthogonally to them. When both gratings had the same orientation, they were also presented in identical phase. In the second interval, a single grating patch was presented on one side of fixation. This grating had the same orientation as the central grating patch presented during the first interval and was always at 30% luminance contrast. We refer to the 30% level of contrast of the second grating presentation as the *standard stimulus*. Following the presentation of the second grating (i.e., the standard stimulus), participants were required to compare the stimuli presented in each interval at the location of the second single (standard) stimulus and indicate with a button press which interval contained the higher contrast.

To direct the allocation of voluntary visual-spatial attention, prior to each trial, one of two possible locations was cued with an arrow appearing at the central fixation. The cue was informative, in that the subsequent judgment was more likely to be performed in that location: the single grating patch in the second interval (i.e., the standard stimulus) appeared on the same side indicated by the arrow on approximately 70% of the trials. Trials in which the standard stimulus appeared in the expected, cued location are henceforth referred to as “validly cued”, or “valid” trials. The 30% of trials in which the standard stimulus appeared in the unexpected, opposite location, are referred to as “invalidly cued” or “invalid” trials.

Previous studies have found a “horizontal effect” in surround suppression, whereby suppression is more pronounced in horizontal orientations, relative to vertical and oblique orientations (Kim, Haun, & Essock, 2010; Kosovicheva et al., 2012). Therefore, in different blocks of trials, participants viewed horizontal or vertical center target gratings and horizontal or vertical surround gratings. On half of the blocks the surround was co-linear and on half of the blocks the surround was orthogonal to the center stimulus. The order of vertical/horizontal stimuli was counter-balanced across subjects, but because the task is more difficult under co-linear conditions, we tested all participants on one of the orthogonal task conditions first (counter-balanced across subjects), to allow them to gradually accommodate to task performance.

In addition to predictive cue blocks, in which an arrow indicated the predicted location of the stimulus in the second interval, subjects were also tested in separate blocks on a version of the task which was identical to the predictive cue condition, except that the cue that appeared before the beginning of each trial did not provide spatial information about the likely location of the target in the trial. In these blocks, the fixation changed into a diamond shape with exactly the same timing to that of the appearance of the arrow in the predictive cue blocks. We refer to this condition as the “neutral cue condition”.

An experimenter monitored eye movements with an infrared eye-tracking camera. Subjects were all experienced psychophysical observers who had previously been extensively trained to fixate while covertly attending to peripheral location, and no eye movements were observed in any of the task performance runs.

### Data analysis

For each level of contrast of the comparison stimulus (i.e., the stimulus that appeared in the first interval, with a surrounding annulus), we calculated the proportion of trials in which the participant responded by indicating that he or she perceived that the comparison stimulus had a higher contrast than the standard stimulus. When the comparison contrast was much higher than the standard stimulus this probability was very high (approaching 1.0) and when the comparison stimulus had a low contrast this probability was very low (approaching 0). The response probabilities for intermediate contrasts follow a sigmoidal shape. To model the regular relationship between contrasts and responses as a continuous function, we considered two models. The first, following Yu et al. (Cong Yu, Klein, & Levi, 2003) was to model this data as a Gaussian cumulative distribution function (cdf). The benefits of this model, apart from the fact that it fits the data well, are that it is a relatively simple model, with only two fit parameters and that these parameters are easily interpretable in terms of the subject’s behavior: One parameter (the mean of the Gaussian distribution) controls the vertical offset of the function and is interpreted as the point of subjective equality (PSE). That is, it is the contrast that the comparison stimulus would need to have in order to appear to have as much contrast as the standard stimulus. This value indicated whether the perception of the comparison stimulus was systematically biased by the presence of the surround. When this term was equal to 0.3 (the displayed contrast) there was no bias and participants were responding to the stimulus veridically. When this number was larger than 0.3 participants were biased to see the comparison stimulus as having lower contrast, even when the contrast was identical. This is an indication of surround suppression due to the presence of the surrounding annulus. The other parameter (the variance of the Gaussian) controls the slope of the function and is interpreted as the sensitivity of subjects to changes in contrast around the PSE. The inverse of the sensitivity is the threshold. That is, when the slope was very high (i.e., high sensitivity), only a small difference in contrasts would be required for participants to detect the presence of a difference in contrasts. If the slope was very shallow, a large difference would be required for participants to reliably detect the presence of a difference in contrasts. Thus, the variance parameter of the Gaussian distribution is an indication of the just noticeable difference (JND).

The other model we considered is the Weibull cdf, following other studies who have used this model to fit psychometric curves to data for discrimination tasks performed with similar stimuli (Kosovicheva et al., 2012; Yoon et al., 2009). This model has 2 additional parameters and in addition to the PSE and JND, this model can explicitly model offsets of the upper and lower asymptotes of the curves, thereby accounting for participants who were overly conservative in their responses and did not consistently respond that the comparison stimulus had higher contrast when this stimulus had much higher contrast than the standard stimulus, or did not consistently respond that the comparison stimulus had lower contrast in trials in which it had a much lower contrast. To determine the model parameters, data for each run was fit to the Gaussian or the Weibull cdf using a variant of the Levenberg-Marquardt algorithm, implemented in scipy.

To compare the models, we used split-half cross-validation (Hastie et al., 2008). This method compares model performance in terms of its accuracy in predicting out-of-sample observations. That is, it tests the ability of a model to generalize to unseen data. Each subject’s data were split into two halves: even trials and odd trials. The models were fit to one half and the parameters were used to predict the performance in the other half: the proportion of “comparison higher than standard” responses in each of the presented contrast values, in each experimental condition. Prediction error was quantified as the root of the mean squared error (RMSE) of the prediction relative to the left-out half (in units of proportion of responses). This procedure was repeated twice, with each half serving both as a ‘training’ set, used to fit the parameters, and as a ‘testing’ set, used to evaluate the model predictions. Lower average cross-validation RMSE in the two iterations indicates better model accuracy, regardless of model complexity (number of parameters).

To determine within-subject reliability of model fits, we performed a boot-strapping analysis: each participant’s data in each condition of the experiment was resampled with replacement to produce 1000 boot-samples, and the models were fit to each of these samples. A confidence interval of the model parameters was computed from the 95 central percentiles of the distribution of parameters across boot-samples for each subject and condition.

To determine the statistical significance of differences between experimental conditions, a mixed-model ANOVA was conducted using R and the ezANOVA package (Lawrence, 2011). Each model parameter was entered into a separate within-subjects ANOVA, with stimulus conditions (horizontal vs. vertical), relative orientation (orthogonal vs. parallel) and attention conditions (valid cue, invalid cue, neutral cue) as factors. Additional ANOVA were conducted with an additional factor of distractor contrast (5 levels).

To promote the reproducibility of our results (Donoho, 2010) and replication by others, all the data and the code used for both stimulus presentation and for data analysis is available to download at http://github.com/arokem/att_ss

**Figure 1:**
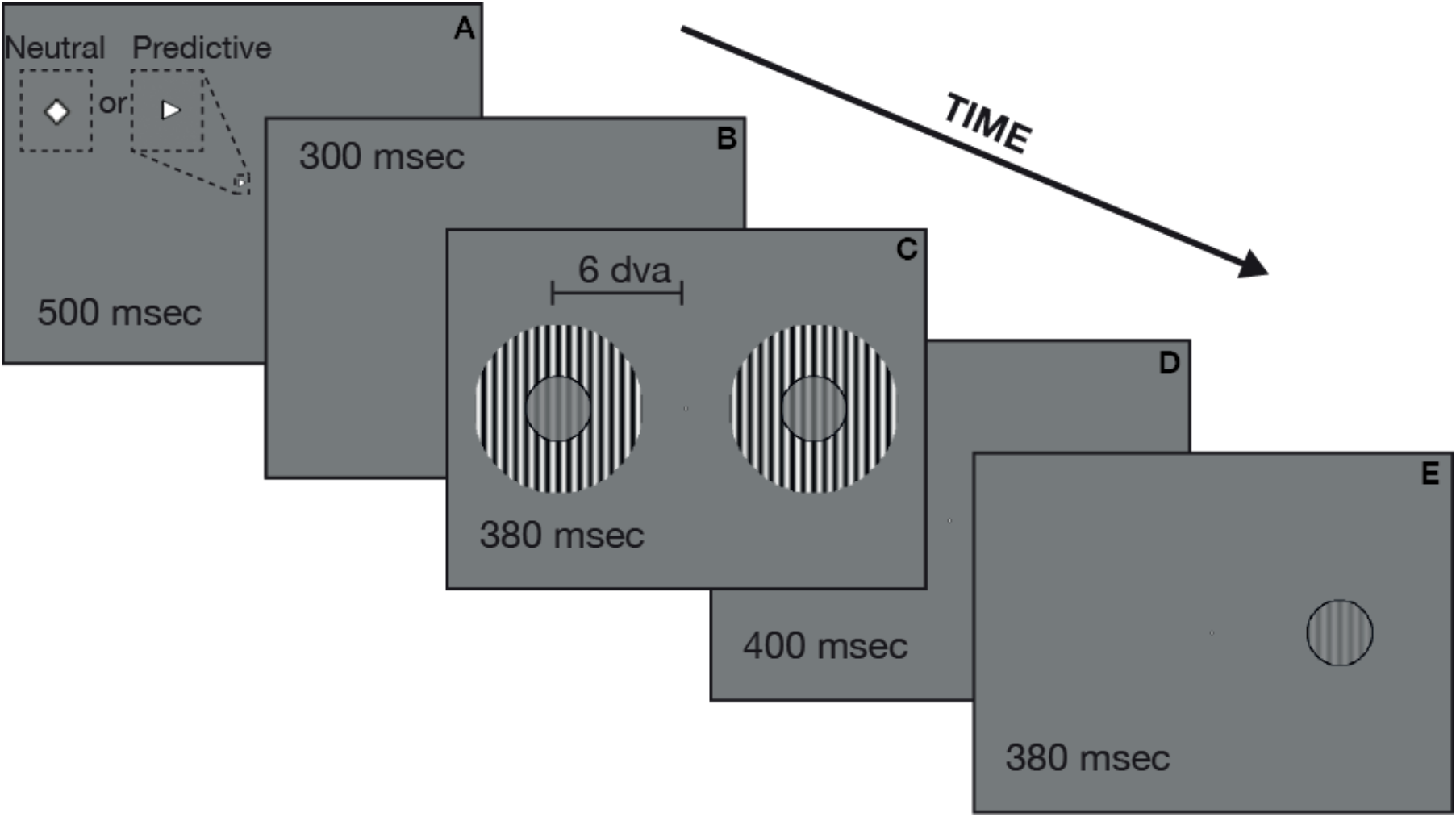
Behavioral paradigm. Orientation-specific surround suppression was assessed using a 2-interval forced choice contrast judgment. At the beginning of each trial a cue was presented (A). In predictive trial cues, the fixation was shaped as an arrow, pointing towards the location in which the contrast judgment was more likely to occur (70% validity). In neutral cue trials, the cue was shaped as a diamond, which provided no information about the location in which the standard stimulus was most likely to occur. Following a cue-to-stimulus interval (B), grating stimuli appeared on both sides of the display (C). We refer to these gratings as the comparison stimulus. These stimuli were surrounded by a co-linear surround (as shown here), or by an orthogonal surround (not shown). After a stimulus-to-stimulus interval (D), a single grating appeared on one side of the display (E). This second stimulus always had 30% contrast and we refer to it as the standard stimulus. Participants were asked to indicate by a button press whether the standard stimulus had higher or lower contrast than the comparison stimulus that appeared in that location in interval C of the trial.

## Results

### Performance in the task can be described using a cumulative Gaussian model

Orientation selective surround suppression (OSSS) was assessed in a contrast-matching task (Figure 1). Participants viewed a grating patch and were asked to indicate whether a preceding comparison stimulus presented in the same location had a higher or lower contrast. Performance can be quantified by asking, for each comparison contrast level, what the probability of responding “comparison higher than standard” was.

For each participant and condition, we fit a cumulative Gaussian function to the data. To validate the choice of this model, we compared the fits of this model to an alternative model: a cumulative four-parameter Weibull function, often used to describe psychometric data in similar tasks (e.g. Kosovicheva et al., 2012; Yoon et al., 2009). To compare the models we use split-half cross-validation (see Methods). We find that the RMSE for the Gaussian model (mean=21.2% ±2%, SEM) is lower than the RMSE for the Weibull model (mean=27.2%±1 SEM), suggesting that the model is a more accurate description of the data, despite the fact that the Weibull model has more parameters. One participant was consistently identified as unreliable based on the bootstrapping confidence intervals of the model fit parameters. This participant had confidence intervals more than 3 standard deviations larger than the group mean in 4 different experimental conditions. Therefore, this participant’s data were excluded from the following presentation of the results. However, the main claims in the text do not change when this participant’s data are included in the analysis. The mean fit of the cumulative Gaussian psychometric curves, not including this participant’s data, is presented in Figure 2.

**Figure 2:**
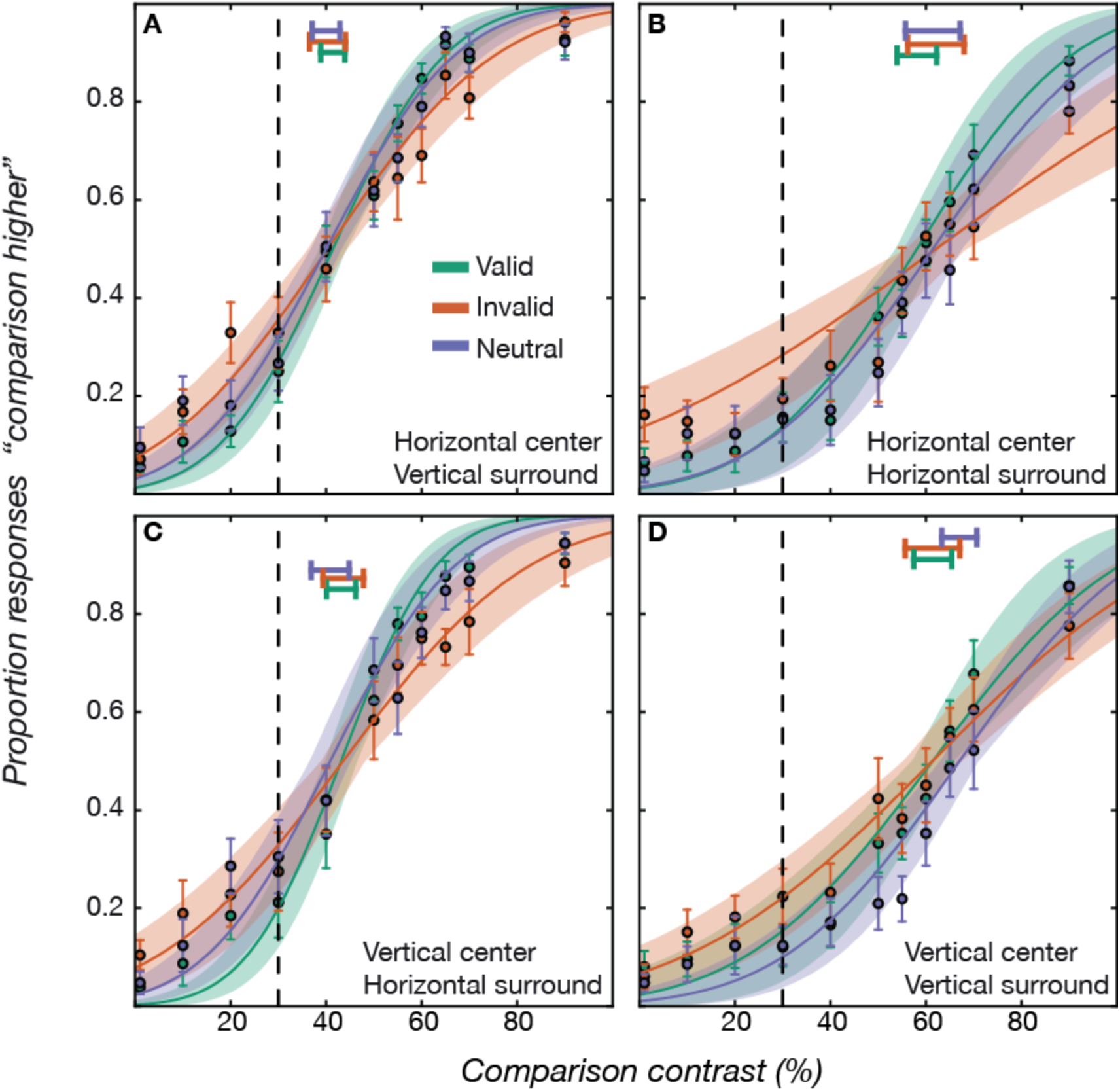
Psychometric curves. Average judgments across subjects are presented (± SEM error bars), for each of the four comparison stimulus conditions: horizontal center (A and B) and vertical center (C and D) by horizontal surround (B and C) and vertical surround (A and D), for each of the attention conditions (valid cue trials: green; invalid cue trials: red; neutral cue trials: blue). Curves depict the cumulative Gaussian model fits (±SEM shaded region). Horizontal bars depict the average bias parameter across subjects (±SEM error bars). Dashed vertical black lines highlight the contrast level of the standard single grating stimulus (30%).

### There is robust OSSS, but there is no effect of focused attention on OSSS

Given that we have chosen to use the Gaussian model, the resulting psychometric curve can be summarized using two parameters of this model: the mean parameter is an indication of the point of subjective equality (or PSE) and the variance parameter is an indication of the just noticeable difference (or JND). The vertical dashed lines in each panel of Figure 2 indicate the veridical contrast of the standard stimulus. If perception was accurate, this is the contrast at which performance on the task should have reached 50%. Instead, the PSE (horizontal bars at the top of each panel indicate the mean ±SEM) is shifted to the right, indicating a bias in the perception of the surrounded comparison grating stimuli in both parallel- and orthogonal-surround stimuli. The comparison stimulus needed to have a contrast higher than the true contrast of the standard stimulus to appear to have the same contrast.

Contrary to previous findings (Kim et al., 2010; Kosovicheva et al., 2012), we did not find a difference in the magnitude of OSSS as a function of absolute orientation of the central grating (no effect of either absolute orientation: F_1,14_=2.76, p=0.12, or interaction of absolute and relative orientations: F_1,14_<1, p=0.8). This means that the two orientations serve as a replication of the same experiment within each subject and serve as an indication of the reliability of the effects we observe here. Nevertheless, following our initial hypothesis, statistical analysis was conducted using absolute orientation as a within-subject factor.

Figure 3 shows each individual’s PSE and group average. The PSE is an indication of the perceived contrast of the stimulus and we find that it is reliably modulated by the relative orientation of the surround: mean PSE is % 32 ± 4.5 (SEM) higher than the comparison contrast for the parallel surround condition and 12% ± 3.2 (SEM) higher than the comparison contrast for the orthogonal surround condition, averaged across attention conditions. This is a replication of the previously well-established orientation-selective surround suppression effect.

On the other hand, within each relative orientation condition, PSE are approximately identical for all the attention conditions and we do not find an effect of attention condition on the PSE (F_1,14_<1, p=0.7), and no interaction between attention condition and relative orientation (F_1,14_=1.4, p=0.25). This suggests that attention does not affect the perceived contrast of the surrounded grating.

**Figure 3:**
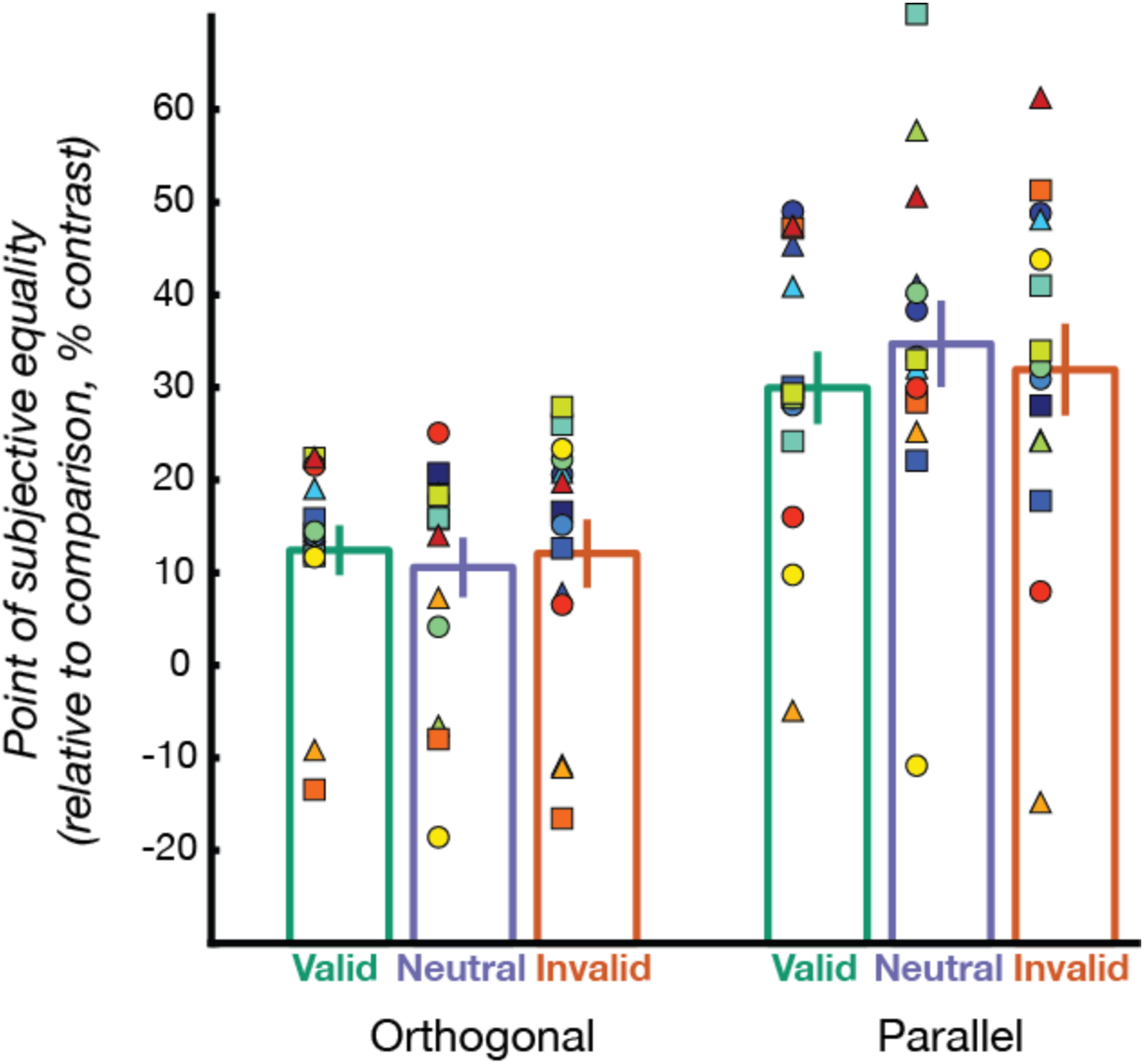
bias. The point of subjective equality (PSE) derived from the fit of the Gaussian model to the data from each subject is presented here. The ordinate axis displays the Gaussian model fit mean parameter, which is an indication of the PSE, or the bias in perception. When the value was equal to 0 perception of the surrounded stimulus was veridical. Values larger than 0 indicate that the surrounded stimulus was perceived as having less contrast indicating suppression. The mean parameter is averaged across presentations of the horizontal and vertical central stimuli in each condition (orthogonal/parallel surround). To compute the PSE relative to the standard stimulus, we subtract 30% from the mean parameter for each condition. Bars denote the group average ± SEM. Filled square/circle/triangle markers denote the performance of individual subjects and each individual subject is identified by a color/shape combination used here and in Figure 4.

### Sensitivity is decreased by both OSSS and diversion of focused attention

The JND is an indication of the sensitivity of the observers to differences in contrast. This can be thought of as the minimal amount of contrast change that has to be introduced in the stimulus in order for participants to perceive that the contrast has changed, and it is a measure of the contrast discrimination threshold at that level of contrast. In the JND parameter, there is a main effect of both relative orientation (F_1,14_=7.7, p<0.05) and attention condition (F_1,14_=6.02, p<0.05). However, there is no interaction between attention condition (cued, neutral, other) and the relative orientation (orthogonal, parallel) in the JND. That is, the effects of attention and surround suppression on threshold are additive. There is only a small difference between the JND in the validly cued trials (orthogonal surround: 17% ± 2 SEM, parallel surround: 30% ± 6 SEM) and JND in the neutral cue trials, which were probed in a separate experimental block (orthogonal surround: 20 % ± 2 SEM, parallel surround: 30 % +/− 6 SEM). Thus, the main effect of attention in the ANOVA is attributed to an effect of the diversion of attention in the focused attention condition, in trials in which the cue was invalid (orthogonal surround: 29.3 % ± 4.8 SEM, parallel surround 43.6 % ± 7 SEM).

**Figure 4:**
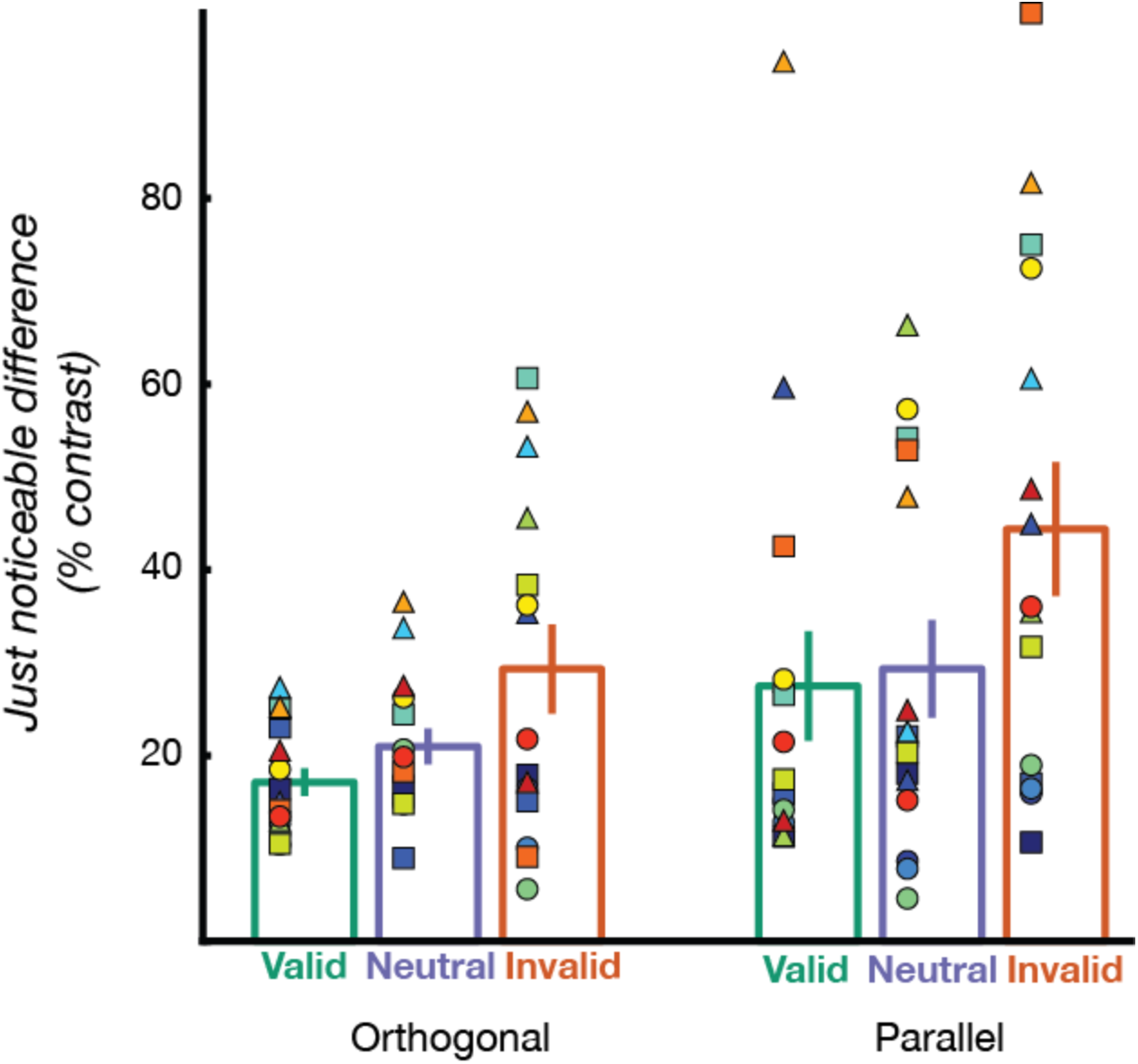
Sensitivity. The just noticeable difference (JND) is derived from the slope parameter of the psychometric curves. The ordinate axis displays the Gaussian model fit variance parameter, which is an indication of the JND, or contrast discrimination threshold. Higher values indicate decreased sensitivity. This is an estimate of the threshold in performing contrast discrimination around the contrast of the standard stimulus. There is a main effect of both surround conditions (JND for parallel higher than orthogonal surround) and of the attention conditions (JND for invalid higher than valid and neutral cue), but no interaction.

### OSSS is modulated by distractor saliency when attention is distributed

In each trial of the experiment, the first interval contained two grating stimuli, one on each side of the screen. While one of these patches ended up being a target for the subsequent discrimination, the other ended up being a distractor, containing no task-pertinent information. To separate the effects of different distractor contrasts in the different cue conditions on bias and sensitivity, we divided the trials in each cue condition according to distractor contrasts. To assure that enough trials were included in the analysis of each distractor contrast, the data (which had 11 contrast levels) were binned into bins of 2 different distractor-contrast levels, covering approximately 10-25% of the contrast scale in each bin, with 3 contrast values in the highest distractor contrast bin. We fit cumulative Gaussian psychometric curves to the data from different distractor contrast bins for each subject, in each attention condition and assessed the effects of distractor contrast on bias and sensitivity (Figure 5).

Replicating the previous analysis of the data, we observed a consistent OSSS across the different distractor contrasts: PSE is always substantially higher for parallel, compared to orthogonal relative-orientations (F_1,14_ =12.5, p<0.001). In addition, in the neutral cue condition (distributed attention), PSE was modulated by distractor contrast in two ways. First, in parallel-surround stimuli, PSE was higher when the distractor had a high contrast than when the distractor had a lower contrast. Thus, suppression was stronger when the distractor contrast was higher. Second, in orthogonal-surround stimuli, PSE was closer to the veridical contrast when the distractor had low contrast, than when the distractor had higher contrast Thus, there was less suppression when distractor contrast was lower. These observations were supported by a significant interaction of cue condition (neutral/valid/invalid) by distractor contrast (F_2,28_=7.42, p<0.01). No other significant effects were found in the ANOVA. Post-hoc within-subjects t-tests revealed that the interaction is mostly driven by a significant difference, in the parallel-surround stimulus, between performance in the highest distractor contrast trials and the lowest distractor contrast trials (t_14_=2.22, p<0.05). No such difference was found in the orthogonal surround conditions (t=1.5, p=0.15), but there were significant differences between the focused attention conditions (average of valid and invalid PSE) and the distributed attention condition (neutral PSE) in the lowest contrast distractor trials in the orthogonal condition (t_14_=2.2, p<0.05) and a near-significant difference between the focused attention condition and the distributed attention condition in the highest contrast distractor trials in the parallel condition (t_14_=1.94, p=0.07).

As observed before, the JND was also significantly affected by relative orientation due to OSSS, with sensitivity substantially decreased in the parallel surround stimuli, relative to orthogonal surround (F_1,14_=11.2, p<0.001). The JND is also affected by cueing condition (F_2,28_=13.1, p<0.001). In the focused attention blocks, the JND is not substantially modulated by distractor contrast in either valid or invalid cue trials. A substantially different pattern appeared in the neutral cue condition: JND in trials in which distractor contrast was low resembled the focused attention validly-cued JND, and JND in trials in which distractor contrast was high resembled JND in the invalidly-cued trials. Accordingly, there is a significant interaction of cue condition by distractor contrast (F_2, 28_====3.7, p<0.05). No other significant effects were found in the ANOVA. Post-hoc t-tests revealed a significant difference between JND in neutral cue blocks and invalid cue trials in the parallel surround condition, when distractor contrasts were low, and this difference is eliminated when distractor contrasts were high (For contrasts smaller than 10%, t_14_=2.6, p<0.05; for contrasts between 10% and 20% t_14_ =2.0, p=0.06; for contrasts between 30% and 40% t_14_=2.9, p<0.05; for contrasts between 50 and 60%, t_14_=0.6, p=0.55; and for contrasts higher than 60%, t_14_= 0.18, p=0.85, Figure 5D). This pattern probably resulted due to the changes in the JND in the neutral cue condition, rather than a tendency of the JND to decrease with increasing distractor contrast in the invalidly cued trials, because this tendency was not reliably observed: the difference between the JND in the invalidly cued trials at the lowest and highest distractor contrasts is not statistically significant (t_14_=1.26, p=0.22; Figure 5D). The complimentary trend appeared in the differences between JND in the neutral-cue condition and the JND in the validly-cued trials, with JND being almost identical in lower distractor contrasts, and diverging at higher distractor contrasts, but the differences that emerged in the high distractor contrasts did not reach statistical significance (For 50-60%: t_14_=1.48, p=0.15; for 65% and above: t_14_=1.42, p=0.18; Figure 5D). A similar pattern appeared in the orthogonal-surround condition: the JND in the neutral condition was similar to the validly cued trials for low distractor contrasts and was similar to the invalidly cued trials at high distractor contrasts. Here, there was no difference between JND of the validly cued trials and neutral blocks in the three contrast bins below 50% contrast (t_14_=0.86,p=0.4; t_14_=0.46,p=0.65; t_14_=1.3,p=0.2; Figure 5C), and a numerical difference that was not statistically significant in the 50-60% contrast bin (t_14_=2.09,p=0.053; Figure 5C). The complimentary pattern was apparent in invalidly cued trials. Here, there was a statistically significant difference between the invalidly cued condition and neutral cue condition for distractor contrasts between 40 and 50% (t_14_=2.4, p<0.05; Figure 5C) and no statistically significant differences in any of the other distractor contrasts. However, we note that the mean JND is higher in the neutral cue blocks than in invalidly cued trials in the two highest distractor contrast bins. This suggests that JND in the neutral-cue condition was similar to the valid cue condition when distractor contrasts were smaller than 50%, and was similar to the invalid cue condition when distractor contrasts exceeded 50%.

Taken together, these patterns of results suggest that when attention is distributed (neutral cue condition) participants engage in a different strategy compared to when attention is focused on one spatial location (valid and invalid cue conditions), even if that location is previously defined as a distractor location (e.g., for invalid trials).

**Figure 5:**
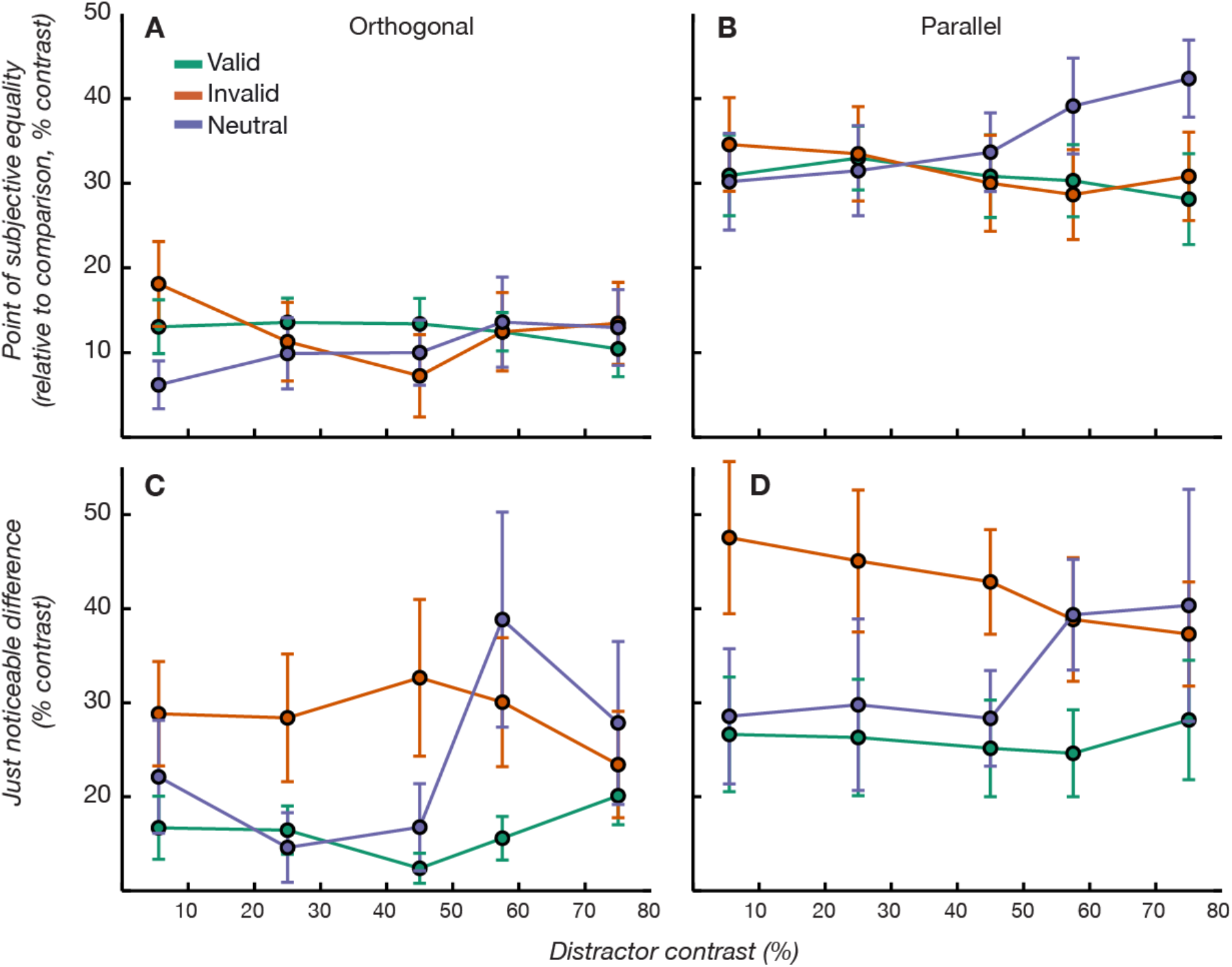
Distractor contrast effects. The contrast of the distractor had very little effect on performance in the valid and invalid cue conditions, but modulated performance in the neutral cue condition. In all these plots, the abscissa denotes the average contrast of the distractor in that bin (for example, the first bin contained data from trials in which the distractor contrast was either 1% or 10%, and the abscissa value is set to 5.5%) **A** In the orthogonal surround, bias decreased for low distractor contrasts. **B** In the parallel surround condition, bias increased for higher distractor contrasts in the neutral cue condition. Sensitivity in the neutral cue condition resembled the sensitivity in the valid cue condition in low distractor contrasts and resembled the sensitivity in the invalid cue condition in high distractor contrasts. This was true in both orthogonal (**C**) and parallel (**D**) surround stimuli.

## Discussion

### Does voluntary attention affect OSSS?

Two measures of performance were used to assess whether attention affects OSSS. The first was the bias induced by OSSS. In the focused attention experiment we found that allocation of voluntary attention to the stimulus, or away from it, does not have a measurable effect on the degree of bias induced by the presence of a surrounding high-contrast stimulus. The contrast at which a suppressed stimulus was judged to have the same contrast as a non-suppressed stimulus was essentially equal in the different attention conditions. This finding seems to be at odds with a line of studies that have found that attention affects the perceived contrast of visual stimuli (Anton-erxleben et al., 2010; Carrasco, Ling, & Read, 2004b; Liu et al., 2009). Importantly, these previous observations were typically made in conditions probing “transient”, or involuntary attention, and not focused voluntary attention (see also Kerzel, Zarian, Gauch, & Buetti, 2010; Prinzmetal, Nwachuku, Bodanski, Blumenfeld, & Shimizu, 1997; Rahnev et al., 2011; Schneider, 2008, 2011). Specifically, some of these studies used relatively short (<250 msec) intervals between the onset of the cue and the onset of the target known to probe involuntary attention, or in tandem with a complex secondary task (Liu et al., 2009). In the present study, we addressed the effects of voluntary attention, at a time-point well after the dissipation of involuntary attention, by setting the cue-to-target interval to a duration of 800 msec (Rokem, Landau, Garg, Prinzmetal, & Silver, 2010). As shown in several previous studies, involuntary attention has different behavioral effects than voluntary attention (e.g. Prinzmetal, Mccool, & Park, 2005; Ling & Carrasco, 2006; for a review see Prinzmetal & Landau, 2007). These two different types of attention also have different neural mechanisms (Esterman et al., 2008; Landau et al., 2007; Rokem et al., 2010). Consistent with those differences, we find that voluntary attention does not alleviate the bias due to surround suppression.

Instead, the most prominent effect of focused voluntary attention on performance was a marked decrease in sensitivity in the trials in which the informative cue was invalid (i.e., misleading), relative to trials in which the informative cue was valid, and trials in which no informative cue was presented (neutral cue condition). While some studies found that neutral cue conditions have an intermediate level of sensitivity, between valid and invalidly cued conditions (Pestilli & Carrasco, 2005; Posner et al., 1980), we found that sensitivity did not substantially differ between the trials in which the cue was valid and the neutral cue condition (when aggregated over all distractor conditions; see further discussion below). However, other studies have shown that that differences in sensitivity between valid and neutral attention conditions can be inconsistent between subjects (Lu & Dosher, 1998; Pestilli & Carrasco, 2007), or depend on the stimulus (Lu & Dosher, 1998) and on the cue type (Jonides & Mack, 1984). In our study, the lack of difference between the valid and the neutral cue conditions is further qualified by the breakdown of these attention conditions by the contrast of the distractor stimulus. This breakdown, which captures differences in the level of distractor saliency in our displays, demonstrates that the perceptual effects of distributed attention differ quite substantially from the effects of focused attention allocated to the stimulus, or allocated away from it.

### OSSS depends on the distribution of attention

We found no interaction between distractor contrast and OSSS in the valid and invalid cue conditions, when the subjects focused their attention on one location. But there was an interaction in the neutral cue condition, when participants did not have information about the location of the task-relevant stimulus, and therefore should have distributed attention between both stimuli. Specifically, bias due to OSSS (PSE) increased in the parallel surround condition when the distractor had high contrast and decreased in the orthogonal surround condition when the distractor contrast was low, bringing perception of that stimulus closer to its veridical contrast. One possible explanation of these results is that the spatial distribution of attention modulated OSSS by integrating more or less of the visual display into the suppressive surround. Thus, contrast perception in the distributed attention experiment was affected by the sum of the contrast on the screen: when the total contrast on the screen was small, bias was reduced (but only in the orthogonal condition). When the distractor contrast was high, the total contrast was larger and bias was increased (but only in the parallel surround condition).

Another explanation is that the changes in sensitivity in the distributed attention experiment were due to capture of involuntary attention to the location of high contrast. Although the timing of the cue-to-target interval was set to allow involuntary effects of the *cue* to dissipate, and to facilitate allocation of voluntary attention, involuntary attention was probably still deployed due to stimulus saliency effects that occur when the two *target* gratings appear. We found that when allocating voluntary attention to a particular stimulus, participants could effectively disregard the information from the uncued location, as the contrast of the stimulus presented in this location had no effect on contrast perception and contrast discrimination in the validly cued trials. This also replicates previous findings showing that distractor contrast has no effect on target contrast gain, when attention is focused (Yigit-Elliott et al., 2011). Moreover, in the invalidly cued trials, there was no effect of the contrast of the stimuli in the cued location, suggesting that processing of the two locations proceeded independently in this condition. When attention is not focused, but distributed over the visual field (neutral cue condition) marked effects of distractor contrast were measured which also depended on surround condition: contrast discrimination thresholds (JND) were lower in the trials in which the distractor had a relatively low contrast (<50%) and resembled the thresholds in the validly cued trials. In trials in which distractor contrast was relatively high (>50%) thresholds increased to resemble those in the invalidly cued condition. Thus, sensitivity in the location of the subsequent target followed a pattern similar to the sensitivity due to allocation or withdrawal of voluntary attention from these locations, in the focused attention condition. This replicates previous findings showing that voluntary attention effects on contrast sensitivity are modulated by the contrast of distractor stimuli in displays with multiple spatially separated stimuli, when attention is distributed (Pestilli et al., 2011; Yigit-Elliott et al., 2011).

Previous studies that examined the interactions of attention and surround suppression used dual-task designs to reduce the availability of attentional resources. As predicted from the classic findings on the effects of inattention on stimulus detection (Bashinski & Bacharach, 1980; Posner et al., 1980), the detectability of a stimulus was reduced when attention was divided between two tasks (Zenger, Braun, & Koch, 1999). However, this effect was much larger when a concurrent mask was presented surrounding the target grating, and in particular when the orientation of the surrounding masks and central target were co-linear. This suggests an interaction between attention and OSSS, where attention is particularly effective in improving detectability when detectability is limited by OSSS. Similarly, surround facilitation effects, which occur in some configurations of co-linear stimuli, are only apparent if the surrounding elements are the target of a secondary task (Freeman, Driver, Sagi, & Zhaoping, 2003; Freeman, Sagi, & Driver, 2001), suggesting again that attention modulates surround interactions. Importantly, because these results were obtained in a dual task design, they were measured under conditions that required distribution of attention over large parts of the visual field, similar to our neutral cue condition. Therefore, in light of these previous studies, we favor an explanation whereby the perceptual bias, and the loss of perceptual sensitivity due to OSSS in our study were indeed modulated by attention, but only when voluntary attention was not specifically allocated to a particular location in the visual field, but was distributed over large portions of the visual field.

While our results relate to previous attempts to explain the neural and perceptual effects of attention in terms of the spatial distribution of attention, and the interactions between different neural pools (e.g. Reynolds & Heeger, 2009; Schwartz & Coen-cagli, 2013), we did not test the predictions of these theories directly, because this would have required a further manipulation of the size of the attentional field (the portion of the visual field to which attention is directed), and this was held fixed in our experiments. Because the surround was never task-relevant, the task design always restricts the attentional field size to the central grating.

### Neural mechanisms of OSSS and attention

Recordings of single cells in early visual cortex show that some cells reduce their response to a stimulus in their classical receptive field (CRF), when an additional stimulus is placed in regions surrounding the RF, and these regions are referred to as the non-classical RF (nCRF) of the cell (Allman, Miezin, & McGuinness, 1985; Hubel & Wiesel, 1965). Co-linear gratings presented in the nCRF modulate responses in the classical RF more strongly (Blakemore & Tobin, 1972; Cavanaugh, Bair, & Movshon, 2002a, 2002b), suggesting that perceptual OSSS depends directly on this form of neural surround inhibition. Non-invasive recordings of human brain activity support this: measurements of surround suppressive effects related to perception have been conducted using fMRI (Kay, Winawer, Rokem, Mezer, & Wandell, 2013; Williams, Singh, & Smith, 2003; Zenger-Landolt & Heeger, 2003), EEG (Kim et al. 2012) and MEG (Haynes, 2003; Ohtani, Okamura, Yoshida, Toyama, & Ejima, 2002). Zenger-Landolt and Heeger (2003) characterized the correspondence between suppression of fMRI BOLD responses and the suppression of perceived contrast finding the best correspondence in the primary visual cortex, V1, rather than visual areas V2 and V3. These ‘higher-order’ areas had a larger degree of suppression than predicted by behavior. This finding suggests that a change in the activity of neural populations in V1 mediates the perceptual effects of surround suppression. Sundberg, Mitchell, & Reynolds (2009) used a display in which stimuli were shown in the nCRF of neurons recorded in visual area V4 of awake macaque monkeys, while the attention of the animal was directed towards or away from a stimulus in the CRF. They found that surround suppression decreased when attention was directed towards the stimulus in the CRF and increased when attention was directed to the stimulus in the nCRF (Sundberg et al., 2009). These results seem to be at odds with our behavioral results, but there are several significant differences. First, there is a difference in the scale of measurement: single neurons vs. behavior. Second, the evidence from fMRI suggests that the locus of perceptual suppression is in V1, rather than higher areas (Zenger-Landolt & Heeger, 2003). Third, there may be important differences between the species that were tested: macaque monkey vs. human. Finally, there are differences in the stimulus design: the stimuli used by Sundberg et al. were two individual separated patches always containing co-linear gratings and they did not directly assess OSSS, instead comparing trials with no surround and trials with a co-linear surround. These differences between the two studies render the comparison somewhat complicated. Still, we cannot reconcile the fact that Sundberg et al. found a roughly 50% decrease in the degree of surround suppression when the center stimulus was attended, relative to when attention was directed away from the stimulus with similar stimulus contrast (33%). We did not find such an interaction between OSSS and attention. Instead OSSS and the allocation of attention to another location seem to sum additively when assessing contrast sensitivity. One possible explanation comes from a study that measured the length tuning of neurons in area V1 (Roberts, Delicato, Herrero, Gieselmann, & Thiele, 2007). Length tuning is also measured by placing stimuli outside the classical RF of a neuron, and is defined as the length of a bar for which a neuron’s response was maximal, after which making the bar any longer caused a decrease in response. In this study, attention was found to decrease length tuning for neurons with RF centered around 2-3 dva eccentricity, and increase length tuning for neurons with RF centered around 6-7 dva eccentricity. This might predict different modulations of OSSS by attention in different eccentricities. In particular, in our study, the stimuli spanned 2-10 dva. Thus, it may be that our lack of attentional modulation simply represents a modulation in one direction in one population of neurons and a modulation in the other direction in other populations. This hypothesis would have to be verified in further experiments in which measurements of neural responses are conducted in tandem with a task design such as the one used here.

## Conclusions

Orientation-selective surround suppression affected both perceived contrast and sensitivity to changes in contrast. On the other hand, the allocation of voluntary visual-spatial attention affected only perceptual sensitivity, but not the appearance of the stimulus. The effects of attention and surround suppression on perceptual sensitivity are independent (combine additively) suggesting that the locus of these effects in the visual system is governed by separate mechanisms. When voluntary attention was focused in one location, other stimuli in the visual field were blocked or attenuated and the contrast of the distractor had no effect on bias, or on sensitivity. When it was distributed over the display, perception could be modulated by the contrast of the distractor: in low distractor contrast conditions, sensitivity was equivalent to the sensitivity observed when voluntary attention was correctly allocated (validly cued). In high contrast distractor conditions, it resembled the sensitivity when voluntary attention was incorrectly allocated (invalidly cued). In addition, perceptual bias due to OSSS was only modulated by contrast in distributed attention conditions, suggesting that when attention is distributed over large parts of the visual field (i.e., in the absence of focused voluntary spatial attention), the triggering of involuntary attention by a salient stimulus can affect the perception of contrast of the eliciting stimulus and of other, less salient stimuli.

## Acknowledgements

The authors would like to thank Franco Pestilli for useful discussions. AR was supported through a fellowship from the National Eye Institute (NEI F32 EY022294) and through a grant from the Gordon & Betty Moore Foundation and the Alfred P. Sloan Foundation to the University of Washington eScience Institute Data Science Environment. AL was supported through a grant from the Israeli Science Foundation and James McDonnell Foundation Scholar Award.

